# Whole-genome methylation profiling of menstrual stem cells identifies novel biomarkers for endometriosis

**DOI:** 10.1101/2025.07.25.666720

**Authors:** Ioanna Tiniakou, Cemsel Bafligil, Raúl Pérez-Moraga, Sarah Harden, Sophie Ribeiro-Volturo, Alfredo Santana Rodríguez, Roberto Notario Manzano, María Alejandra Santana Suárez, Marta Tortajada Valle, María Ángeles Martínez-Zamora, María Teresa Pérez Zaballos, Alicia Martin Martinez, Francisco Carmona, Cristina Fernández-Molina

## Abstract

Endometriosis, despite its high prevalence, is underdiagnosed and poorly managed due to lack of clinically validated biomarkers and pathophysiological insight. Menstrual blood-derived stem cells (MenSCs) have been implicated in disease pathogenesis, but their diagnostic potential remains unexplored. We conducted a clinical study (n=42; 19 endometriosis, 23 controls) to assess whether DNA methylation profiles of freshly isolated MenSCs can identify disease-specific biomarkers. Whole-genome methylation sequencing revealed differentially methylated regions (DMRs) enriched in genes linked to hallmarks of endometriosis (e.g., inflammation, tissue remodelling, development). These DMRs robustly distinguished cases from controls, independent of technical and clinical variables. Machine learning models trained and validated on these DMRs achieved high diagnostic performance (specificity 83%, sensitivity 79%). Integration with an independent single-cell RNA sequencing dataset showed that the DMRs may modulate gene expression, further supporting their biological relevance. These findings position MenSC DNA methylation profiling as a promising, non-invasive approach for early endometriosis diagnosis and personalised care.

## Introduction

Endometriosis is a common gynaecological disorder characterised by extra-uterine growth of endometrial-like tissue and chronic low-grade inflammation^1^. Despite an estimated prevalence of 10% among women of reproductive age, diagnosis is typically delayed by an average of 7 to 10 years^2^. This stems from a limited understanding of disease pathogenesis and lack of reliable non-invasive diagnostics, often leading to misdiagnosis^3^. Endometriosis presents with diverse non-specific symptoms that may not reflect disease severity, rendering clinical assessment and symptom management challenging. It often causes debilitating pelvic pain, fatigue, and infertility, significantly impairing quality of life and mental health. Historically, the gold standard for diagnosis has been laparoscopic surgery, during which endometriotic tissue biopsies are obtained for histological confirmation. To reduce diagnostic delay and facilitate early intervention, recent guidelines from the European Society of Human Reproduction and Embryology recommend a combined assessment of clinical and imaging findings to establish a diagnosis^4^. These guidelines strongly advise against the use of biomarkers for diagnostic purposes, citing insufficient clinical validation and reproducibility. Post-diagnosis, endometriosis management remains complex as existing staging systems offer limited prognostic value and poorly correlate with symptom severity or treatment response. Molecular biomarkers may address these gaps by advancing the understanding of disease mechanisms, enabling earlier detection, and identifying disease subtypes linked to symptomatology and therapeutic response.

Menstrual blood has emerged as a valuable, non-invasive source of stem cells with mesenchymal properties, commonly referred to as menstrual blood-derived stem cells (MenSCs)^5^. When localised in the perivascular regions of the basalis and functionalis layers of the endometrium, these cells are termed as endometrial mesenchymal stem cells (eMSCs) and contribute to endometrial regeneration and repair during the menstrual cycle^6^. MenSCs exhibit high proliferative and migratory capacity, robust colony-forming ability and the potential for multilineage differentiation^7^. These features, combined with their endometrial origin, have positioned MenSCs at the centre of several theories on endometriosis pathogenesis.

Retrograde menstruation, the leading proposed mechanism underlying endometriosis, suggests that viable endometrial cells, including MenSCs, reflux through the fallopian tubes into the pelvic cavity where they can implant, grow and form endometriotic lesions^8^. Although retrograde menstruation is common (approximately 90% of women)^9^, only a subset of individuals develop endometriosis, suggesting differences in MenSCs may contribute to disease susceptibility^10^. The stem cell theory further proposes that MenSCs can disseminate through lymphatic or haematogenous routes resulting in lesions at both pelvic and extra-pelvic sites^11^. The high proliferative and migratory potential of MenSCs, demonstrated both *in vitro* and *in vivo*, lend support to this mechanism^12,13^.

MenSCs express mesenchymal stem cell markers such as CD105, CD90, CD73, CD44 and CD29^14,15^ that are also preserved in eMSCs found in endometriotic lesions^16^. In endometriosis patients, MenSCs show dysregulation of genes associated with proliferation, immune modulation, and hormone responsiveness^17^. These findings suggest that MenSCs contribute to both lesion initiation and disease maintenance, underscoring their diagnostic and therapeutic potential.

To date, studies attempting to characterize MenSCs have predominantly utilised cultured cells. However, even short-term culture is known to alter the epigenetic and transcriptional landscapes of cells, restricting the relevance of such findings to primary cells^18^. In contrast, direct isolation from menstrual blood allows for non-invasive, phase-standardised sampling of MenSCs, circumventing the need for cell culture. This approach ensures high specificity by targeting a single disease-relevant population, thereby enhancing molecular resolution and enabling precise analysis of native omic states in both health and disease. Among these, DNA methylation stands out for its chemical stability, resistance to short-term physiological fluctuations, and growing application in *in vitro* diagnostic devices^19^. In endometriosis, altered methylation has been linked to modified gene expression, leading to increased cellular proliferation, invasion, and progesterone resistance^20^.

Here, we investigate whether methylomic profiling of MenSCs can identify biomarkers for endometriosis. We performed enzymatic methylation sequencing (EM-seq) on fluorescence-activated cell sorting (FACS)-isolated MenSCs from women with or without endometriosis and identified differentially methylated regions (DMRs) linked to endometriosis-related pathways. The identified DMRs were then evaluated as non-invasive biomarkers using machine learning models trained on MenSC-derived methylation data. Finally, we explored the functional relevance of these methylation patterns by integrating them with publicly available single-cell RNA-seq (scRNA-seq) data from eutopic endometrium, revealing transcriptional changes associated with the disease. Taken together, our findings provide insights into the epigenetic landscape of endometriosis and highlight MenSCs as a promising source of novel disease-specific biomarkers.

## Results

### Endometriosis and control samples exhibit similar metrics and processing outcomes

Participant recruitment was conducted across two clinical centres and sample processing was performed as outlined in Fig. 1A (for details see “Methods” section). A total of 42 participants, including 19 diagnosed with endometriosis and 23 non-endometriosis controls (for details see “Methods” section), were included in the analysis. Participants in the control and endometriosis groups had mean body mass indexes (BMIs) of 21.55 and 21.94, mean ages of 33.04 and 37.11 years, mean ages of menarche of 12.57 and 12.05 years, and mean durations of menstrual bleeding of 5.00 and 4.79 days, respectively (Table 1). Menstrual blood volumes and flow rates were similar between the two study groups (median volume: 5.0 vs 5.0 mL; median flow rate: 0.50 vs 0.54 mL/h, control vs endometriosis), with no significant differences detected (Fig. 1B).

**Figure 1.**
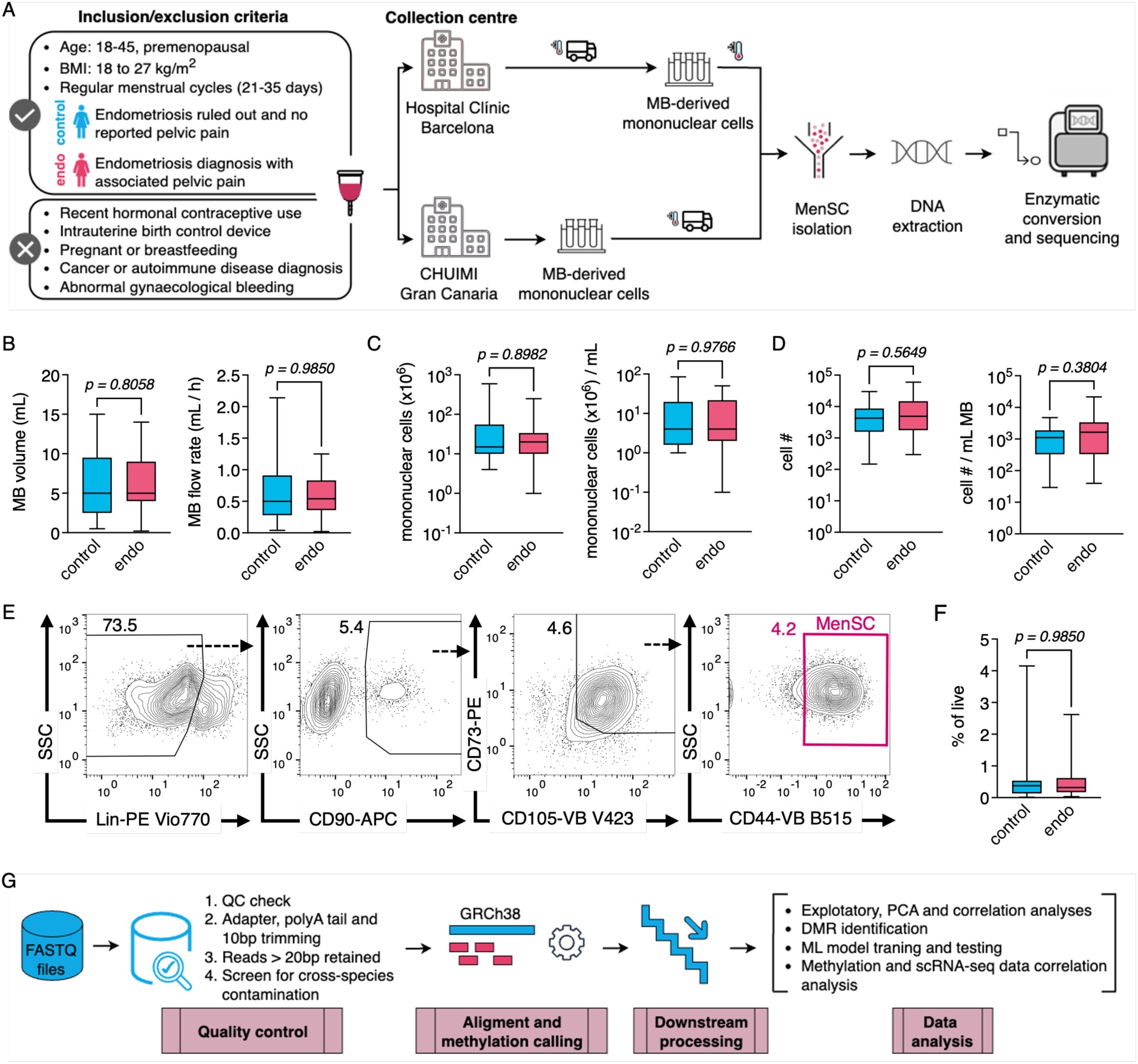
Experimental workflow and quantitative profiling of MenSC isolation from menstrual blood. **A** Schematic of participant recruitment and sample processing workflow. **B** Menstrual blood volume and flow rate collected overnight among participants of the different collection centres. Shown are box plots of menstrual blood volume or volume per hour. **C-D** Absolute numbers of **C** Ficoll-isolated mononuclear cells and **D** sorted MenSCs. Shown are box plots of total cell numbers and cell numbers per mL of collected menstrual blood. **E** Phenotypic profile of MenSCs (Lin-CD90+ CD73+ CD105+ CD44+) within the mononuclear cell population in menstrual blood. Shown is representative staining plot corresponding to the gating strategy used to isolate the cells by FACS. Cells were gated on viable cells. Numbers represent frequencies among total live cells. Lineage was defined as CD45+ CD34+ CD14+ CD19+ HLA-DR+. **F** Frequency of sorted MenSCs among total live cells. **G** Schematic of bioinformatic analysis workflow. Box plots included values of n=42 participants. Whiskers in box plots indicate minimum and maximum values. Significance was determined using the Mann-Whitney test. *P*-values are as indicated in the respective graphs. *BMI* body mass index, *endo* endometriosis group, *MB* menstrual blood, *MenSC* menstrual-blood derived stem cell, *PCA* principal component analysis, *DMR* differentially methylated region, *scRNA-seq* single-cell RNA sequencing, *ML* machine learning.

**Table 1.**
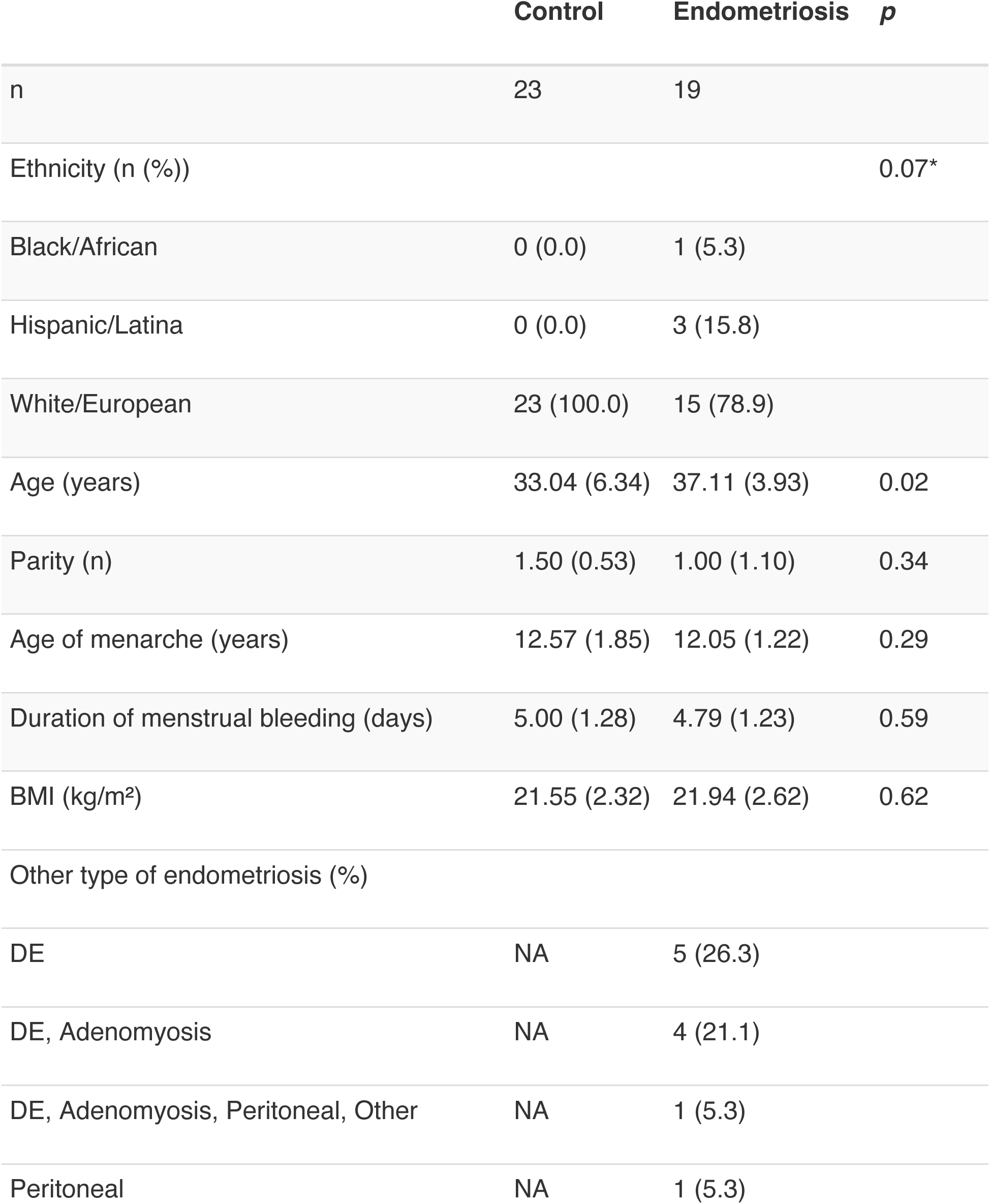
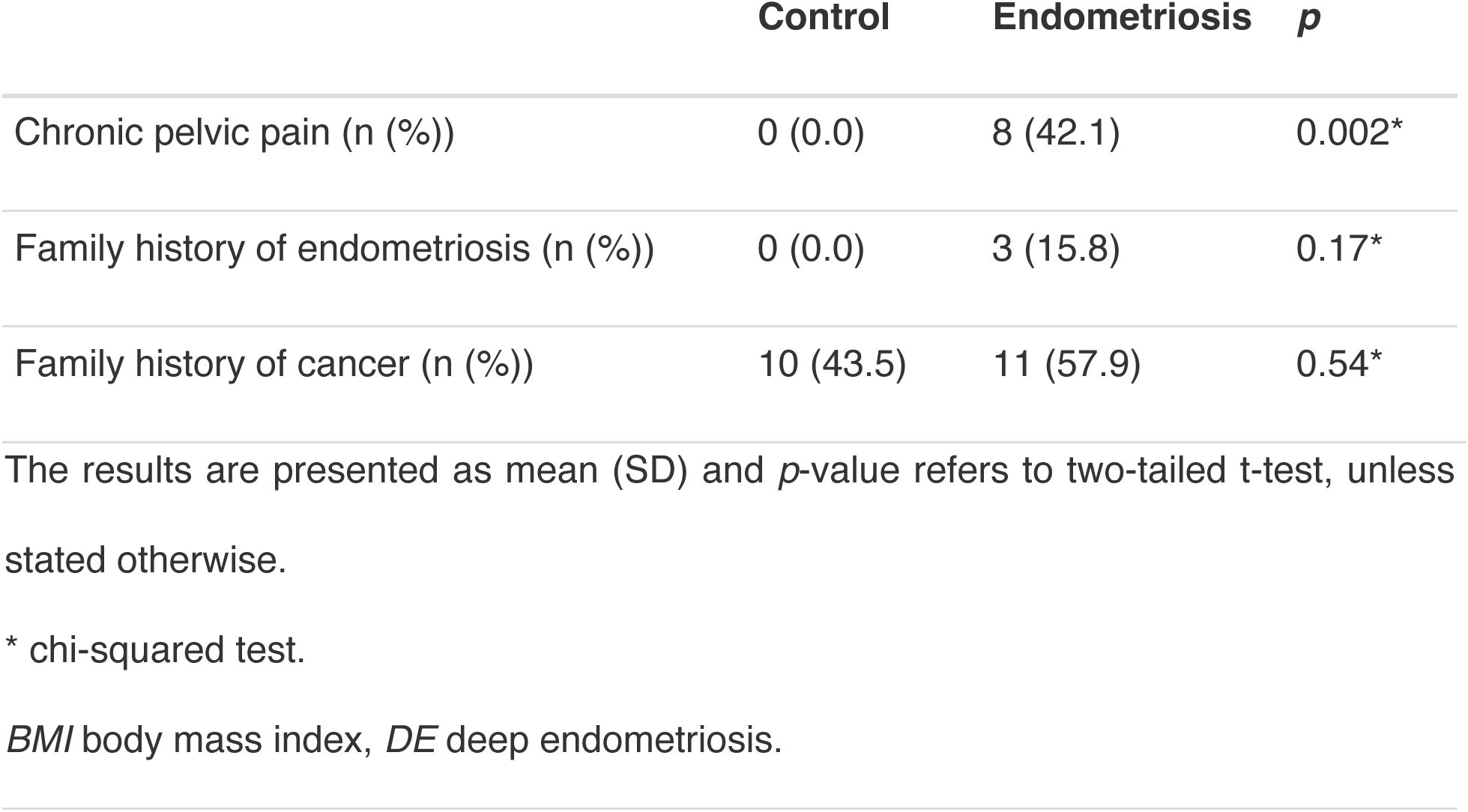
Demographic and clinical characteristics of study participants.

To isolate MenSCs from menstrual blood we performed a two-step procedure involving mononuclear cell isolation followed by sorting of the target cells. Quantitative analysis revealed no significant differences in mononuclear cell yield (median total cell count: 15 x 10^6^ vs 20 x 10^6^; median cell count per mL of MB: 4 x 10^6^ vs 4 x 10^6^, control vs endometriosis) or in the number of sorted MenSCs (median total cell count: 4293 vs 4904; median cell count per mL of MB: 1112 vs 1639, control vs endometriosis) between the two groups (Fig. 1C, D). To achieve selective isolation, we characterised MenSCs using a combination of multiple negative selection markers along with four established positive markers specific to this cell type (CD90, CD73, CD105, CD44; see details in “Methods”; Fig. 1E). No significant differences were observed in the frequency of MenSCs among live cells between the two groups (median frequency: 0.37 vs 0.32 %, control vs endometriosis; Fig. 1F). EM-seq of sorted MenSCs from each sample yielded methylation profiles, which were subsequently analysed using several downstream approaches (Fig. 1G). These included exploratory data analysis, identification of an endometriosis-associated DMR signature, development of a machine learning-based diagnostic model, and independent validation of the DMR signature’s impact on the eutopic endometrium transcriptome.

### DMR analysis reveals predominant hypermethylation in MenSCs in endometriosis and distinct separation of clinical groups

To account for potential confounding effects in downstream analysis, we first assessed baseline clinical variables. Age was the only variable to differ significantly between control and endometriosis groups (*p*=0.02; see Table 1). Therefore, age was included as a covariate in the generalised least squares model used for identifying DMRs. Differential methylation analysis at the region level identified a total of 466 DMRs, with 458 regions being hypermethylated and 8 regions hypomethylated (imputed *p* < 0.05), indicating a strong MenSC hypermethylation profile in endometriosis patients. This marked overrepresentation of hypermethylated regions aligns with the overall methylation distribution observed in CpG positions (Supplementary Fig. 1). Notably, the hypermethylation patterns observed in endometriosis exhibit parallels with those documented in other conditions such as ovarian cancer^21,22^, uterine fibroids^23^, and other cancers^24^.

In terms of genomic context, the majority of identified DMRs (362 DMRs) were located in open sea CpG regions, with the remainder distributed across CpG shores (42 DMRs), CpG shelves (38 DMRs), and CpG islands (24 DMRs) (Fig. 2A). Genomic annotation indicated that most DMRs were located within gene bodies (238 intronic and 56 exonic regions), while a substantial proportion (70 DMRs) was associated with transcriptional regulatory elements, including promoter regions.

**Figure 2.**
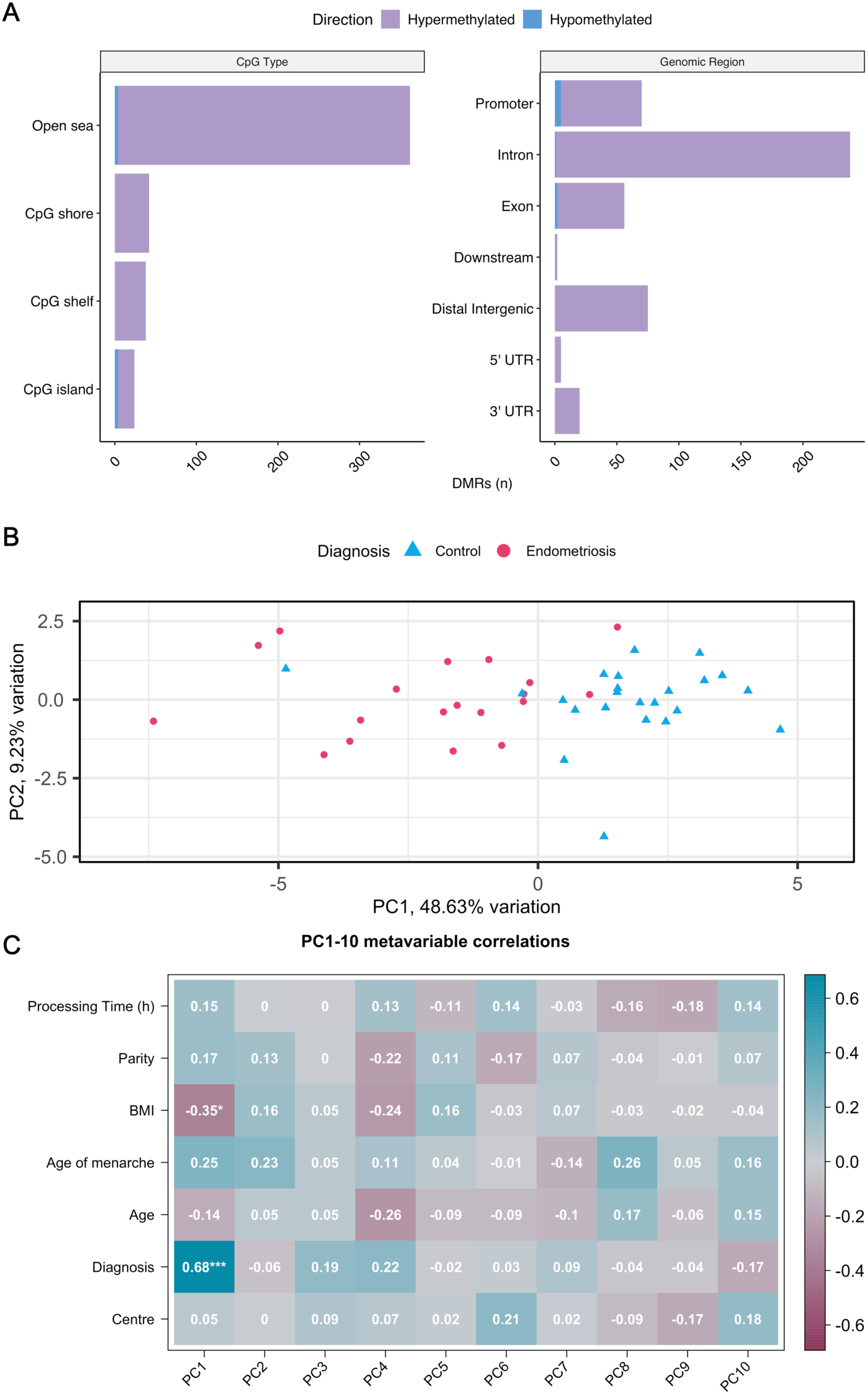
Exploratory analysis of EM-seq data and DMR detection. **A** Genomic context of the detected DMRs, showing CpG annotations and genomic region types. **B** Principal Component Analysis (PCA) plot of the 19 DMRs selected by the feature-selection procedure common to both Random Forest (RF) and Support Vector Machine (SVM) algorithms. **C** Correlation plot depicting relationships between the selected DMRs and various clinical variables (Pearson correlation *p*-value thresholds: 0.05 < *p* < 0.01 = *, 0.01 < *p* < 0.001 = **, *p* < 0.001 = ***).

To identify the most critical DMRs that effectively distinguish between endometriosis and control samples, a dual feature selection approach was employed, combining Support Vector Machine (SVM) and Random Forest (RF) algorithms. This approach yielded a consensus list of 19 key DMRs identified as critical for distinguishing endometriosis from controls. To assess their discriminative capacity, an unsupervised PCA was performed, revealing distinct methylation patterns differences between the two groups (Fig. 2B). Principal component analysis (PCA) results showed that PC1 accounted for 48.63% of the total variance and distinguished endometriosis and control samples, supporting further exploration to refine candidate biomarkers.

To further evaluate the relevance of the selected DMRs, we examined the correlation between PCs and the clinical and technical variables included in the study. The top 10 PCs were analysed, as they collectively accounted for 93.5% of the total variance in the dataset. Figure 2C demonstrates that PC1 showed a strong correlation with the diagnosis status (correlation coefficient = 0.68*, p* < 0.005) and a weak correlation with BMI (correlation coefficient = 0.35, *p* < 0.05). No significant correlations were observed with the remaining clinical variables analysed, including processing time and collection centre, highlighting the robustness of the protocol and resilience to pre-analytical variability. Taken together, these results indicate that PC1 captures critical endometriosis-related biological variability, highlighting the promise of the selected DMRs for further evaluation.

### DMR signature highlights genes with disease-relevant functions and pathways

To assess the discriminative power of the identified 19-DMR signature, we performed hierarchical clustering using Euclidean distance. The DMR-based clustering accurately classified the majority of endometriosis and control samples (Fig. 3A) supporting the robustness of the signature in distinguishing endometriosis samples from non-endometriosis controls.

**Figure 3.**
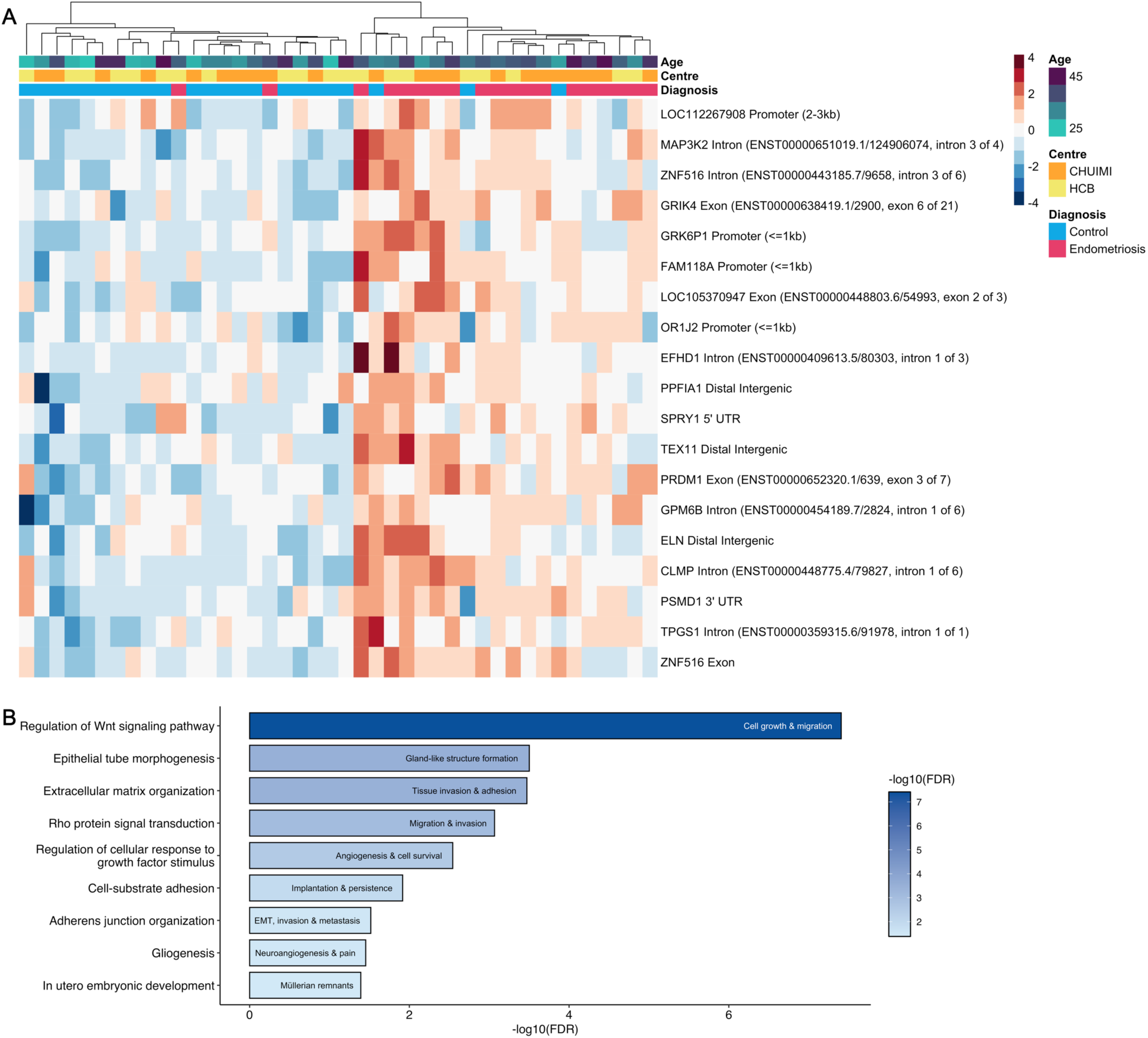
Comprehensive analysis of DMRs and their functional annotation**. A** Heatmap displaying the 19-DMR signature identified through the dual feature selection algorithm. Red tiles represent hypermethylated DMRs, while blue tiles indicate hypomethylated DMRs. **B** Summary of the ORA analysis for GO biological processes. Bar labels represent biological processes associated with the respective GO terms relevant to endometriosis. *ORA* over-representation analysis, *GO* gene ontology

Key genes captured in the DMR signature included *ZNF516*, *GPM6B, PSMD1 and CLMP.* Notably, the dual feature selection algorithm identified multiple DMRs within intronic and exonic regions of *ZNF516*, a DNA-binding protein which regulates transcriptional activity of downstream target genes. Previous studies have reported *ZNF516* to be hypermethylated and transcriptionally downregulated in endometriomas^25^. *GPM6B*, a membrane glycoprotein involved in cell-cell communication, was reported to be downregulated in eutopic endometrial samples in a multi-omics study^26^. Moreover, loss of *GPM6B* expression has been linked to the development of a mesenchymal subtype in glioblastoma^27^. These findings suggest that GPM6B may play a critical role in the progression of various pathological conditions, including endometriosis. A hypermethylated region within the 3′ UTR of *PSMD1,* a regulatory subunit of the 26S proteasome, may interfere with proteasomal function and potentially contribute to the abnormal tissue growth and inflammation observed in endometriosis^28^. Furthermore, Psmd1 dysregulation has previously been reported in mouse xenograft models of endometriotic lesions, suggesting a direct link between its epigenetic regulation and disease pathogenesis. Lastly, DMRs mapped to the first intron of *CLMP,* a transmembrane protein localised at junctions between endothelial and epithelial cells, could lead to aberrant cell-cell adhesion and disruption of endometrial stromal cell integrity (Human Protein Atlas)^29^. Furthermore, CLMP is a known tumour suppressor in colorectal cancer, where its loss has been associated with WNT/β-catenin pathway activation and enhanced cellular proliferation^30^. These observations highlight the role of CLMP dysregulation in endometriosis and potentially other proliferative disorders^31^.

To characterise the identified DMR signature and its potential impact at functional level, Gene Ontology (GO) over-representation analysis was performed using the 458 hypermethylated DMRs described above. As shown in Fig. 3B, several endometriosis-relevant biological processes were found to be significantly enriched. Notably, multiple terms related to the WNT signalling pathway emerged among the most overrepresented categories (Supplementary Fig. 2). Dysregulation of WNT signalling is linked to aberrant cellular proliferation, invasion, and tissue remodelling in endometriosis^32^. In addition, enrichment of terms related to extracellular matrix (ECM) organisation was observed. Genes within the DMR signature likely contribute to ECM dynamics and adhesion properties, potentially promoting the invasive behaviour of MenSCs^33,34^. Moreover, the involvement of angiogenesis, highlighted by terms such as “Regulation of cellular response to vascular endothelial growth factor stimulus” further supports the notion that neovascularisation is critical to lesion establishment and maintenance^35,36^. Finally, several processes associated with epithelial and tissue morphogenesis were listed among the enriched terms^37^. These results imply that the epigenetic alterations identified in MenSCs could be compatible with cellular behaviours that support the establishment and maintenance of endometriotic lesions, such as tissue remodelling, cellular invasion and adhesion, and the initiation of neovascularisation. Collectively, our findings highlight the biological relevance of the genes within the MenSC-derived DMR signature, linking significant DNA methylation changes to key pathways associated with endometriosis pathogenesis.

### DMR signature-based machine learning model accurately diagnoses endometriosis and outperforms prior approaches

To evaluate the predictive capacity of methylation profiling of MenSCs, we trained and tested multiple machine learning algorithms to distinguish between endometriosis and control samples. Given the limited sample size, a leave-one-out (LOO) validation pipeline was implemented to minimize overfitting and provide a realistic estimate of diagnostic performance (Supplementary Fig. 3).

We trained 14 different models with varying numbers of DMRs (ranging from 2 to 20), ranked by their imputed *p*-values in each training iteration. Most models achieved a mean accuracy exceeding 0.90 in the training phase, except for LightGBM and LightGBMXT models, which maintained accuracies of ∼0.50 (Fig. 4A). During the testing phase, the top three performers; Weighted Ensemble L2, ExtraTreesGini, and NeuralNetTorch, achieved accuracies of 0.81, 0.76, and 0.76, respectively, using four DMRs (Fig. 4B; Supplementary Fig. 4). Focusing on the best-performing Weighted Ensemble L2 model, we calculated an accuracy of 0.81, a specificity of 0.83, and a sensitivity of 0.79 (Fig. 4C). Consistent with our previous exploratory analysis, the most influential DMRs driving classification (Fig. 4D) were those located in the exonic and intronic regions of *ZNF516*, as well as in distal intergenic and promoter regions of *LOC101929614* and *LOC112267908*, respectively. These findings reinforce the diagnostic potential of leveraging methylome dysregulation in MenSCs to accurately detect endometriosis in a non-invasive manner.

**Figure 4.**
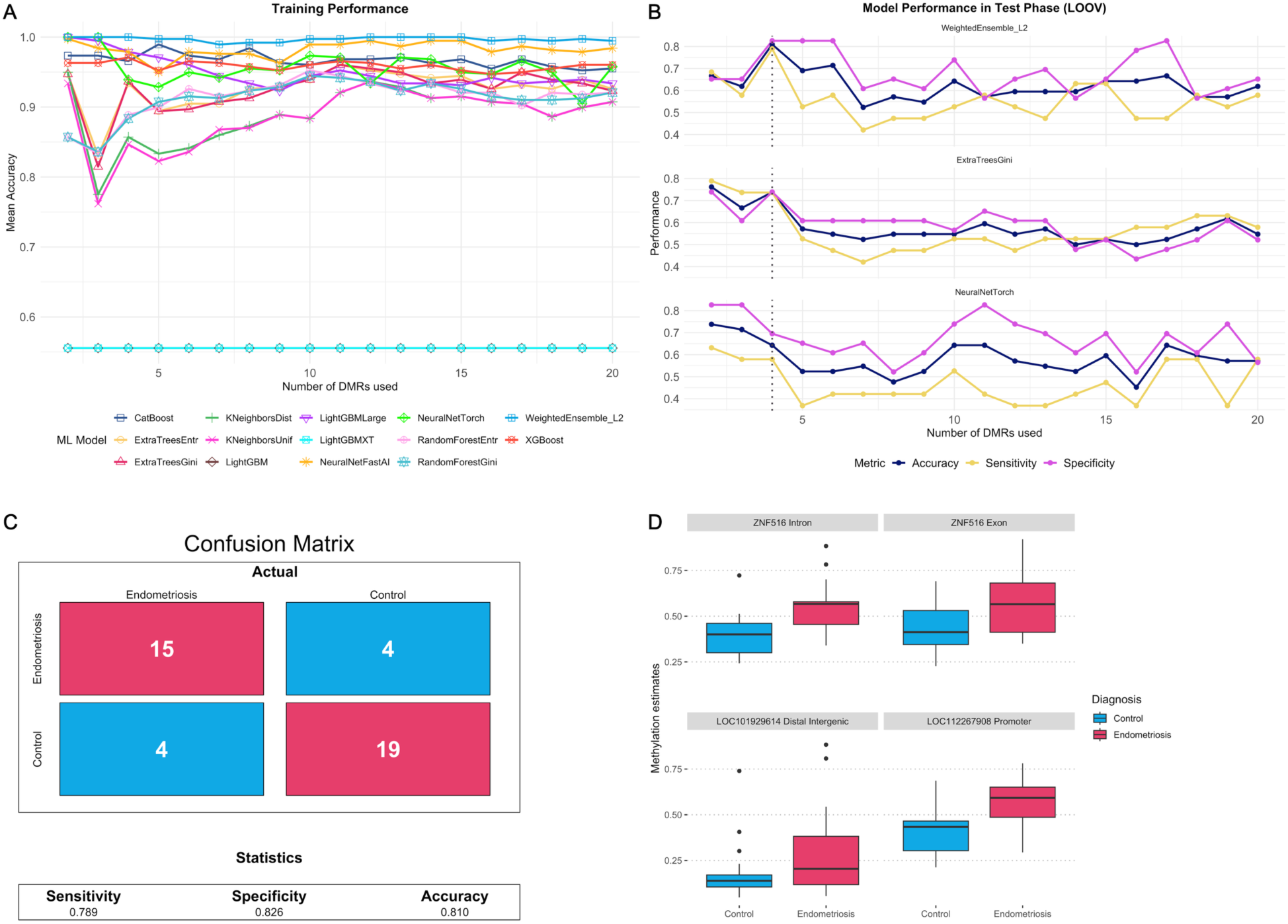
Machine learning model performance using a leave-one-out training and testing pipeline. **A** Line plot showing the mean accuracy during the training stage, evaluated using between 2 to 20 DMRs. **B** Line plot illustrating the accuracy, sensitivity, and specificity of the top 3 best-performing models in the test stage. The dotted line indicates the optimal number of DMRs to use, which is 4. **C** Confusion matrix displaying the performance of the best model, WeightedEnsemble L2. **D** The methylation levels of the top 4 DMRs utilized by WeightedEnsemble L2.

### MenSC-derived methylation signature is associated with suppressed target gene expression in an independent eutopic endometrium scRNA-seq dataset

To independently validate the effect of the identified DMR signature on the endometrium, we used the Human Endometrial Cell Atlas (HECA) dataset^38^, the most comprehensive scRNA-seq dataset of eutopic endometrium from individuals with and without endometriosis. Figure 5A depicts the Uniform Manifold Approximation and Projection (UMAP) plot of 159,052 mesenchymal lineage cells, comprising perivascular, smooth muscle, and stromal cell populations. These cells were sampled across various stages of the menstrual cycle, from the early proliferative phase to the late decidual stage, thereby capturing the complete transcriptomic landscape of stromal cells throughout the cycle in both experimental groups (Supplementary Fig. 5). To detect the possible presence of stem cell-like cells, we calculated a module score using MenSC markers to identify potential stromal stem cells within the eutopic endometrium. By applying a module score cutoff of ≥ 0.8, we detected 1,198 cells (Fig. 5B), which were locally enriched in early-proliferative and proliferative stromal (eStromal) cell clusters. Most of these cells mapped to the eStromal (stromal cells in early-proliferative and proliferative phases) and eStromal_MMP (stromal cells in the menstrual phase) clusters (Fig. 5C). To investigate the transcriptomic differences in stromal stem cells between endometriosis and control samples, we selected cells annotated as eStromal or eStromal_MMP. This yielded a total of 826 cells (403 endometriosis and 423 control cells).

**Figure 5.**
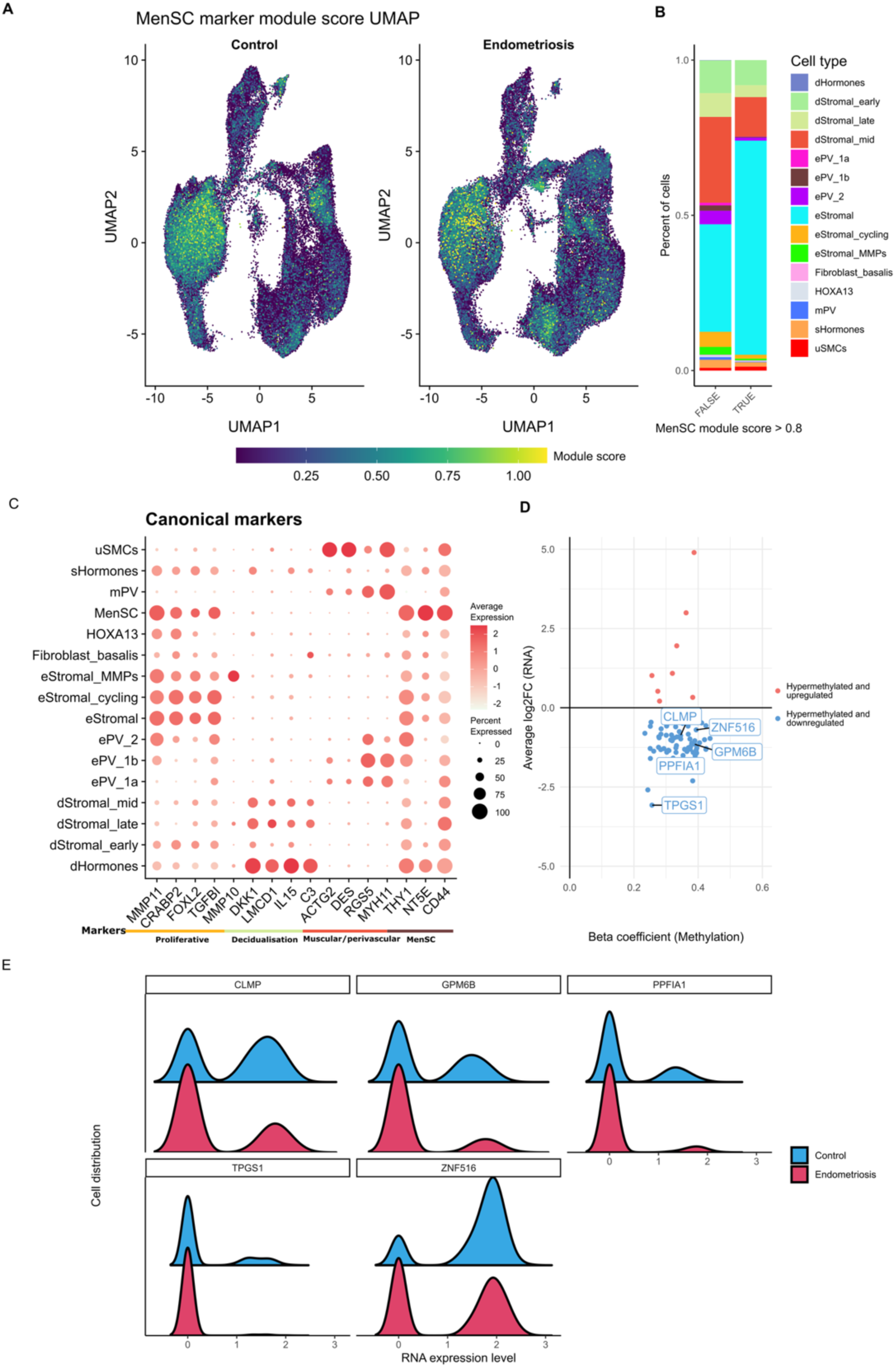
Integrative analysis of the DMR signature detected in the HECA scRNA-seq external dataset. **A** UMAP plot displaying the module score for MenSC markers. **B** Bar plot showing the distribution of cell types between cells with a MenSC module score > 0.8 and all remaining cells. This reveals an enrichment of stromal compartment cells in proliferative phase. dHormones, hormone-induced decidualised stromal cells; dStromal_early, decidualised stromal cells (early stage); dStromal_late, decidualised stromal cells (late stage); dStromal_mid, decidualised stromal cells (mid stage); ePV_1a, endometrial perivascular cells; ePV_1b, endometrial perivascular cells; ePV_2, endometrial perivascular cells; eStromal, endometrial stromal cells (proliferative phase); eStromal_cycling, endometrial stromal cells in cycling stage; eStromal_MMPs, endometrial stromal cells (menstrual/early proliferative phase); Fibroblast_basalis, fibroblast cells from the basalis part; HOXA13, HOXA13 positive cells; mPV, myometrial perivascular cells; sHormones, hormone-exposed stromal cells; uSMCs, uterine smooth muscle cells. **C** Dot plot displaying the markers of the cellular subpopulation. **D** Scatter plot of the beta coefficient versus the average log2 fold change. Each point represents a DMR associated with a gene present in the scRNA-seq dataset and identified as differentially expressed (FDR < 0.05) in stromal cells labelled as eStromal and eStromal_MMPs. The highlighted points are genes detected using the dual machine learning feature selection strategy from the previous analysis. **E** Ridge plot illustrating the expression of these selected genes in the aforementioned stromal cells (normalized log-transformed expression values). *FDR* false discovery rate, *UMAP* Uniform Manifold Approximation and Projection.

We next performed differential gene expression (DGE) analysis comparing endometriosis to controls cells and correlated these results with our previously characterised DMR signature. A substantial proportion (∼90%) of genes that overlap both hypermethylated DMRs and DEGs showed reduced expression in endometrial stromal stem cell populations from endometriosis-affected individuals (Fig. 5D). These findings support a strong correlation between the MenSC methylation signature and the transcriptional profiles observed in the MenSC marker-positive cells in the HECA dataset. Interestingly, several genes identified through our dual feature-selection algorithm (*ZNF516*, *TPGS1*, *GPM6B*, *CLMP*, and *PPFIA1*) were downregulated at the transcriptional level (Fig. 5E). The results shown here suggest that the identified methylation signature has a functional impact at the transcriptome level of MenSC marker-positive cells within the eutopic endometrium of patients.

## Discussion

Menstrual blood provides a non-invasive window into the uterine environment, with low intra-cycle biological variability, making it suitable for molecular profiling of endometriosis, and other gynaecological diseases. In this study, we validated the robustness of our protocol for DNA methylation analysis in MenSCs and identified disease-specific methylation signatures with strong diagnostic performance, achieving 81% accuracy in testing. We further demonstrated the biological relevance of the identified DMRs through external transcriptomic correlation and pathway analysis, which implicated processes involved in disease development and persistence. Altogether, these findings support the potential of menstrual blood-derived epigenetic biomarkers to advance our understanding of endometriosis and improve its detection.

Previous studies on methylation changes in endometriosis have largely used array-based methods which are prone to batch effects^39^ and cover only about 1.5-3% of all CpG sites^40^. These studies were performed on bulk tissue, potentially obscuring cell type-specific methylation patterns in heterogeneous tissues such as the endometrium. Notably, changes in rare cell populations such as eMSCs, which comprise only 0.02% to 1.23% of endometrial cells, may be masked in such analyses^41^. This challenge is further compounded by the fact that the endometrium undergoes substantial methylation changes throughout the menstrual cycle, an effect not observed in peripheral blood^42^. A recent study by Mortlock et al., analysing samples from 984 participants, provided foundational insights into the impact of menstrual cycle phase on endometrial methylation. The study showed that methylation changes were predominantly driven by cycle phase, with disease-specific signals emerging only in moderate-to-severe cases (i.e., stage III and IV combined)^43^. Our study design overcomes previous limitations by isolating MenSCs directly from menstrual blood and performing cell type-specific whole genome methylome profiling on primary, freshly isolated cells for the first time. The collection of menstrual blood standardised the sample timing across participants, minimising phase-dependent variability. Profiling freshly isolated MenSCs allowed an unbiased, comprehensive analysis of the epigenetic landscape in this disease-relevant key cell population, facilitating direct comparisons between patients and controls. Finally, the application of a novel whole genome methylation sequencing protocol that avoids harsh bisulfite treatment, and therefore better preserves DNA integrity and improves coverage uniformity, enabled accurate, high-resolution profiling of CpG methylation in MenSCs. MenSCs have been previously implicated in the aetiopathogenesis of endometriosis, as discussed earlier^44^. Building on this, our findings demonstrate that the genome-wide methylation profile of these cells reflects disease-specific alterations, providing molecular evidence for their role in ectopic lesion development. In support, GO analysis revealed methylation changes in genes associated with key mechanisms of endometriosis pathogenesis and lesion persistence. These included regulation of the WNT signalling pathway, indicating epigenetic modulation of key signalling processes. Additionally, epigenetic dysregulation of pathways governing stem cell activity may further contribute to ectopic lesion initiation and progression.

Our study demonstrates that a machine learning-based approach leveraging DMRs from isolated MenSCs achieves a diagnostic accuracy of 0.81, with a specificity of 0.82 and a sensitivity of 0.79. This performance is particularly notable given the current state of biomarker-based diagnostic tools for endometriosis, which, despite extensive research, have yet to produce a clinically validated, non-invasive diagnostic method^44,45^. To date over 1,100 potential biomarkers, primarily proteins and RNAs, have been identified across various biological compartments, including peripheral blood, peritoneal fluid, follicular fluid and urine. However, only four of these candidates (TNF-α, MMP-9, TIMP-1, and miR-451) have been consistently validated across multiple studies^45^. Even among these, sensitivity and specificity vary substantially depending on patient phenotype, menstrual cycle phase, and comorbidities, underscoring the complexity and heterogeneity of endometriosis^45,46^. In contrast, our model exhibited robust predictive performance using four DMRs, highlighting the potential of methylation-based classification as a more reliable approach to non-invasive endometriosis diagnosis. Moreover, our findings support the use of menstrual blood as a stable diagnostic sample, circumventing the biological variability associated with menstrual phase-sensitive biomarkers and leveraging epigenomic signatures which are inherently more stable than miRNAs or proteins^47^. Nevertheless, to fully capture the clinical and molecular heterogeneity of endometriosis and enhance clinical applicability of this approach, validation in larger, more diverse patient cohorts is essential.

The identified MenSC-derived DMR signature not only distinguishes endometriosis samples at the epigenetic level but also demonstrates potential functional relevance through its influence on gene expression as supported by analysis of an independent transcriptomics dataset. Specifically, analysis of HECA scRNA-seq dataset revealed that genes marked by hypermethylation in our DMR signature were significantly downregulated in eutopic stromal stem cell populations from endometriosis patients. This result indicates that the detected epigenomic signature may influence transcription changes.

This study has a few limitations that should be considered. First, the low yield of sorted MenSCs resulted in limited DNA quantities obtained, which constrained sequencing depth and reduced the extent of methylation changes that could be detected across the genome. This limitation reflects the biological challenge of working directly with a rare cell population without prior *in vitro* expansion. Second, the relatively small sample size reduces the statistical power of the study and limits the ability to perform stratified analysis. Nevertheless, the use of whole-genome methylation sequencing provides high-resolution, unbiased coverage across the genome, enhancing our capacity to detect meaningful disease-associated methylation changes despite sample size constraints. To further mitigate the impact of limited sample size, we adjusted the statistical cutoff parameters accordingly and emphasise the importance of independent validation in larger, multi-ancestry cohorts. Third, although our study controls were selected based on absence of clinical suspicion or history of endometriosis, and confirmed by negative imaging or surgical findings, the inherent limitations of current diagnostic methods mean that undiagnosed disease cannot be entirely ruled out. This common challenge in endometriosis research may lead to conservative estimates of test specificity. Finally, while we assessed the transcriptional impact of the DMR signature using scRNA-seq data from eutopic endometrium, these data were not derived from the same MenSC population. In the absence of transcriptomic data from freshly isolated MenSCs, we focused on stromal cells expressing canonical MenSC markers to approximate the *in vivo* population following menstrual shedding. The observed downregulation of hypermethylated genes in these cells supports the functional relevance of our methylomic signature across molecular layers and biological contexts and underscores the need for direct transcriptomic profiling of freshly isolated MenSCs in future studies.

In conclusion, our findings highlight the potential of MenSCs to reveal endometriosis-associated molecular alterations. The identified methylation signatures were not only predictive but also mapped to pathways implicated in lesion formation, including WNT signalling, stem cell regulation, and extracellular matrix remodelling. Their transcriptional suppression in MenSC-like stromal populations further supports a functional role in disease pathogenesis. By providing a stable, cell-type-specific molecular readout, this approach offers a promising avenue for non-invasive diagnosis and, with further validation, may inform future efforts toward more individualised patient care. Larger and more diverse studies will be essential to assess clinical utility across varying disease presentations.

## Materials and Methods

### Ethics statement

The study was approved by the Ethics Committee for Investigation with medicinal products (CEIm) of both hospitals: the Ethics Committee for Clinical Research of Hospital Clínic de Barcelona (reference number: HCB/2023/0050) and the Ethics Committee of Las Palmas H.U.G.C. Dr. Negrín (reference number: 2022-435-1). Written informed consent was obtained from all participants. All experiments were conducted in compliance with the ICH E6 Good Clinical Practice guidelines and the Declaration of Helsinki.

### Participant Recruitment

Participants were recruited through the Hospital Clínic de Barcelona (Barcelona, Spain) and the Complejo Hospitalario Universitario Insular Materno Infantil (CHUIMI; Gran Canaria, Spain) according to the following inclusion criteria: (i) premenopausal women aged 18 to 45 years; (ii) BMI of 18-27 kg/m^2^; (iii) history of spontaneous regular menstrual cycles (21-35 days); (iv) no menstrual cycle-altering drug treatment (e.g. oral contraceptive pills) for at least 3 months prior to participation. The number of patients recruited was similar across the two centres. Participants in the endometriosis group had a confirmed diagnosis of ovarian endometriosis, identified by laparoscopy and/or imaging within the previous year, in accordance with the ESHRE endometriosis diagnosis guidelines^4^. While ovarian endometriosis served as a minimum inclusion criterion, many participants presented with additional lesion types (see Table 1). Eligible endometriosis participants reported pelvic pain symptoms commonly associated with the condition, such as dysmenorrhea, dyspareunia, and dyschezia. Non-endometriosis participants had no history or suspicion of endometriosis, had endometriosis ruled out by the same diagnostic methods, and reported no pelvic pain or infertility issues. All participants were further excluded if (i) pregnant or breastfeeding; (ii) used an intrauterine birth control device; (iii) experienced abnormal gynaecological bleeding without a known cause; (iv) had history of severe cardiac, respiratory, renal, endocrine or haematological diseases; or (v) were diagnosed with any type of cancer and/or autoimmune disease.

### Menstrual blood sample collection and initial processing

Participants collected menstrual blood overnight on the first or second day of their cycle using menstrual cups. Upon collection, menstrual blood was immediately transferred into sterile 50 mL conical tubes containing 30 mL collection medium (DMEM/F12, 1% Pen/Strep 100 U/mL, 1% Amphotericin B 100X, 1% L-Glutamine 2 mM, EDTA 2 mM) and maintained at 4°C until further processing. Menstrual blood samples collected in Hospital Clínic de Barcelona were shipped at 4°C and processed at endogene.bio’s laboratories within 48 hours. Upon receipt, sample temperature and volume were measured, and a rapid HIV test (ReLab) was performed. Samples collected in CHUIMI were processed in the hospital’s laboratories using the same standardised protocol. The isolated mononuclear cells were frozen and shipped to endogene.bio’s laboratories for further processing.

### Mononuclear cell isolation by density gradient centrifugation

Mononuclear cells were isolated from collected samples by centrifugation on a Ficoll gradient. Briefly, menstrual blood in collection medium was washed with 1x PBS (Gibco) and filtered to remove blood clots and tissue. The filtered suspension was slowly overlaid on Ficoll-Paque PREMIUM (Cytiva) and mononuclear cells were isolated by density gradient centrifugation^48^.

### Flow cytometry and MenSC isolation by FACS

Menstrual blood derived-mononuclear cells prepared as described above were washed and counted on a MACSQuant Analyzer 10 (Miltenyi Biotec). Cells were subsequently stained for multicolour analysis and sorting. Lineage (Lin) cocktail included antibodies to human CD45, CD34, CD14, CD19 and HLA-DR. Fluorochrome-conjugated antibodies against human CD90, CD73, CD105 and CD44 were used to identify the MenSC population. Viobility 405/520 Fixable dye was used to assess cell viability, and staining was performed in 1x PBS for 15 min at RT in the dark. Staining for extracellular markers was performed in FACS buffer (1x PBS, 0.5% BSA, 2 mM EDTA) for 20 min at 4°C in the dark. Samples were acquired on MACSQuant Analyzer 10 (Miltenyi Biotec) and further analysed with FlowJo v10.10.0 software (Tree Star). Cell sorting was performed on the MACSQuant Tyto Cell Sorter (Miltenyi Biotec). Statistical analysis of flow cytometry data was performed using Prism (GraphPad). Normal distribution of data was not assumed, and statistical significance of differences between experimental groups was determined by non-parametric Mann-Whitney test.

### Sample preparation for EM-seq

Genomic DNA was isolated using NucleoSpin Tissue XS or NucleoSpin Tissue Mini kit (Macherey-Nagel) and quantified on a Qubit 4 Fluorometer using the Qubit dsDNA High Sensitivity Assay Kit. DNA quantities ranging from 0.5 ng to 10 ng were sheared to ∼350-550 bp length by Covaris LE220 Plus and used as input to generate sequencing libraries. The libraries were prepared and amplified using the NEBNext Enzymatic Methyl-seq Kit (New England Biolabs) according to manufacturer’s instructions, adjusting the PCR cycles based on starting DNA quantity. The sequencing libraries passing technical QC were sequenced in 150 bp paired-end mode on Illumina NovaSeq X Plus using 6-8% unmethylated PhiX Control v3 (Illumina).

### EM-seq Data Quality Control

Fastq files were pre-processed to assess data quality and identify potential sequencing issues. Briefly, the quality of reads was assessed using FastQC^49^, followed by adapter, 10 bp low-quality ends and polyA tail trimming with TrimGalore^50^. Reads with less than 20 bp after trimming were discarded. To investigate cross-species contamination, FastQ Screen was used to perform mapping of samples with a high percentage of duplicates (> 60%)^51^.

### EM-seq Data Read Alignment and Methylation Calling

Trimmed Reads were mapped to the human GRCh38 reference genome using the default parameters of BSBolt (v1.6.0), which integrates a modified BWA-MEM algorithm optimised for both bisulfite and enzymatic methylation sequencing data^52^. Post-alignment, duplicate reads were removed using samtools (v1.21)^53^. Methylation calling was performed using BSBolt with default settings, generating output in CGmap format. These files were then converted to Bismark-compatible CpG report format using a custom Python script to facilitate downstream analysis.

### EM-seq Data Downstream Analysis

The total number of CpGs analysed was assessed using the *comethyl* (v1.3.0) package^54^ to explore various coverage thresholds and shared CpG positions across samples. Coverage thresholds ranging from 1x to 10x per CpG, and the proportion of samples required for a CpG to be considered valid, were evaluated. Final cutoffs were set at a minimum coverage of 3x in at least 80% of samples. After applying these cutoffs, *dmrseq* ^55^ and *bsseq* ^56^ (implemented via *DMRichR* (v1.7.8)^57^) were used to identify and calculate DMRs in the dataset. The discovery process utilised a smoothing and weighting algorithm to adjust CpG counts based on coverage. Candidate background regions were defined by grouping CpGs with similar genomic proximity and methylation levels. Statistical significance of the regions was determined through permutation testing, with empirical *p*-values calculated by comparing observed test statistics against a null distribution generated from 41 permutations. Analysis parameters included a minimum CpG coverage of 3x, at least five CpGs per DMR present in 80% of samples, and a single-CpG coefficient cutoff corresponding to a 6.5% methylation difference, with adjustment for age. Finally, DMRs were annotated using a combination of the *annotatr* (v1.32.0)^58^ and *ChIPseeker* (v1.42.1)^59^ R packages. Promoter regions were first defined using the *build_annotations* function from *annotatr* with the hg38 genome build. DMRs were then annotated to genomic features (e.g., promoters, introns, exons) using the *annotate_regions* function from the *DMRichR* pipeline, incorporating genome-wide annotation tracks. To assign each DMR to the nearest gene, we used *annotatePeak* from *ChIPseeker* with the TxDb.Hsapiens.UCSC.hg38.knownGene and org.Hs.eg.db databases. The resulting annotations included gene symbols, gene names, and functional context relative to transcription start sites (TSS).

### Selection of Informative DMRs

From the identified DMRs, the *methylLearn* function was used to select the most informative regions distinguishing between endometriosis and control samples. This function applies RF and SVM algorithms to identify key DMRs. We selected the top 5% of DMRs that were consistently prioritised by both algorithms. To evaluate these DMRs, PCA was performed using the *PCAtools* package^60^. Data were scaled and centred with the *pca* function, and correlations between DMRs and various clinical or experimental variables were assessed using *eigencorplot*.

### Enrichment Analysis

Overrepresentation analysis (ORA) was performed using the *WebGestaltR* (0.4.6) package^61^, considering only genes corresponding to hypermethylated DMRs. GO Biological Process terms were selected as the ontological framework, with minimum and maximum category sizes set to 10 and 500, respectively. All genes annotated in the hg38 genome assembly were used as the reference set, and genes identified in the DMR signature served as the input list. Statistical significance was determined using false discovery rate (FDR) adjusted *p*-values, with a cutoff of 0.05.

### Machine learning

To prevent potential data leakage into the training set and ensure reliable test results given the limited dataset size, an LOO data split strategy was implemented. The dataset was divided into 42 folds, with each fold consisting of 41 samples for training and 1 independent sample for testing. DMR detection was performed on the training data using the same settings described in the “EM-seq Data Downstream Analysis” section. The resulting CpG matrices were smoothed, and features were derived from the top 2 to 20 DMRs, ranked by imputed *p*-value. These reduced matrices were then used to train machine learning models using the Python library AutoGluon^62^ with the “medium_quality” setting. The following models were trained: LightGBM, CatBoost, XGBoost, Neural Networks (Torch and FastAI), Random Forest (Gini and Entropy), Extra Trees (Gini and Entropy), Weighted Ensemble, and k-Nearest Neighbors (kNN). Accuracy was used as the primary metric during training. For each fold and each number of DMRs used, predictions on the test sample (excluded from training) were collected to evaluate overall model performance in the test phase. Final performance metrics included sensitivity, specificity, and accuracy.

### Methylation and scRNA-seq validation data

The impact of the DMRs identified in this study on the transcriptome was investigated using the state-of-the-art scRNA-seq dataset of eutopic endometrium from the HECA dataset^38^. This dataset comprises 313,527 cells from 63 women, both with and without endometriosis, integrated from multiple studies. First, the mesenchymal lineage was extracted and re-clustered using the Seurat R package (5.1.0). The count matrix was normalised by dividing each cell’s feature counts by the total counts for that cell, multiplying by a scale factor, and then applying a natural log transformation using the *log1p* function. Next, the most variable RNA features were identified and the top 2,000 were selected for integration using the *SCTransform* function. Data integration was performed using the *harmony* (1.2.3) package and integrated by “dataset” variable to remove batch effects related to dataset origin. PCA was performed and the top 30 PCs were used to compute the UMAP. The *FindNeighbors* and *FindClusters* functions were applied for graph-based clustering by constructing a kNN graph using Euclidean distance in the PCA space. Clusters were subsequently defined using the Louvain algorithm to optimise the standard modularity function.

To detect the stromal stem cell population within the mesenchymal lineage, the *AddModuleScore* was used with previously established MenSC marker genes (*THY1*, *NT5E*, *CD44*). This function calculates the average expression of a specified gene set per cell, subtracting the aggregated expression of control feature sets. All analysed features are binned based on average expression, and control features are randomly selected from each bin. Log-normalised counts were used for this analysis. A cut-off of ≥ 0.8 was used to select highly enriched cells expressing MenSC-associated markers^63^. Finally, the stromal cells from the proliferative phase were exclusively selected to perform downstream analysis. DGE analysis was performed on the selected cells using the Wilcoxon test, with *p*-values corrected by the FDR. Finally, the DMR and DGE results were integrated to assess the correlation between the DMR signature identified in MenSCs and in a stromal stem cell population from an external eutopic endometrium dataset.

## Acknowledgements

We thank Life & Brain GmbH for providing sequencing services. We are grateful to the physicians who contributed to the clinical study: Meritxell Gracia, Georgina Feixas, Victoria Sánchez Sánchez, Patricia Esther Escamilla, Luciana Obreros, Mariazel Pérez, Neuda Marqués de Oliveira, José Vázquez Nuñez, Patricia Hernández Delgado, and Juan José Artazkoz Marques de Oliveira. We also acknowledge CRAnarias, Investigación y Desarrollo S.L for clinical study management, and Teresa Galera Monge for coordinating the study internally. We thank Nilufer Rahmioglu and Altuna Akalin for their scientific support, and Pablo Arriagada and Verónica Alam for their clinical and operational guidance.

## Author Contributions

I.T., S.H., and S.R.V. performed the experiments. I.T., C.B., and R.P.M. analysed the data and interpreted the results. R.N.M. conducted the literature review and provided scientific support for the project’s development. M.T.P.Z. conceived the project and secured funding. C.F.M. led the overall project design and supervision. A.M.M. and F.C.H. acted as clinical principal investigators. A.S.R., M.A.S.S., M.Á.M.Z., and M.T.V. collected and processed clinical samples. I.T., C.B., R.P.M., and C.F.M. wrote the manuscript with input from all authors.

## Competing Interests Statement

C.B., R.P.M., I.T., S.H., S.R.V., R.N.M. and C.F.M. are employees of endogene.bio. M.T.P.Z. is the Chief Executive Officer of endogene.bio. All other authors declare no competing interests.

