## Supplementary figures for "Whole-genome methylation profiling of menstrual stem cells identifies novel biomarkers for endometriosis"

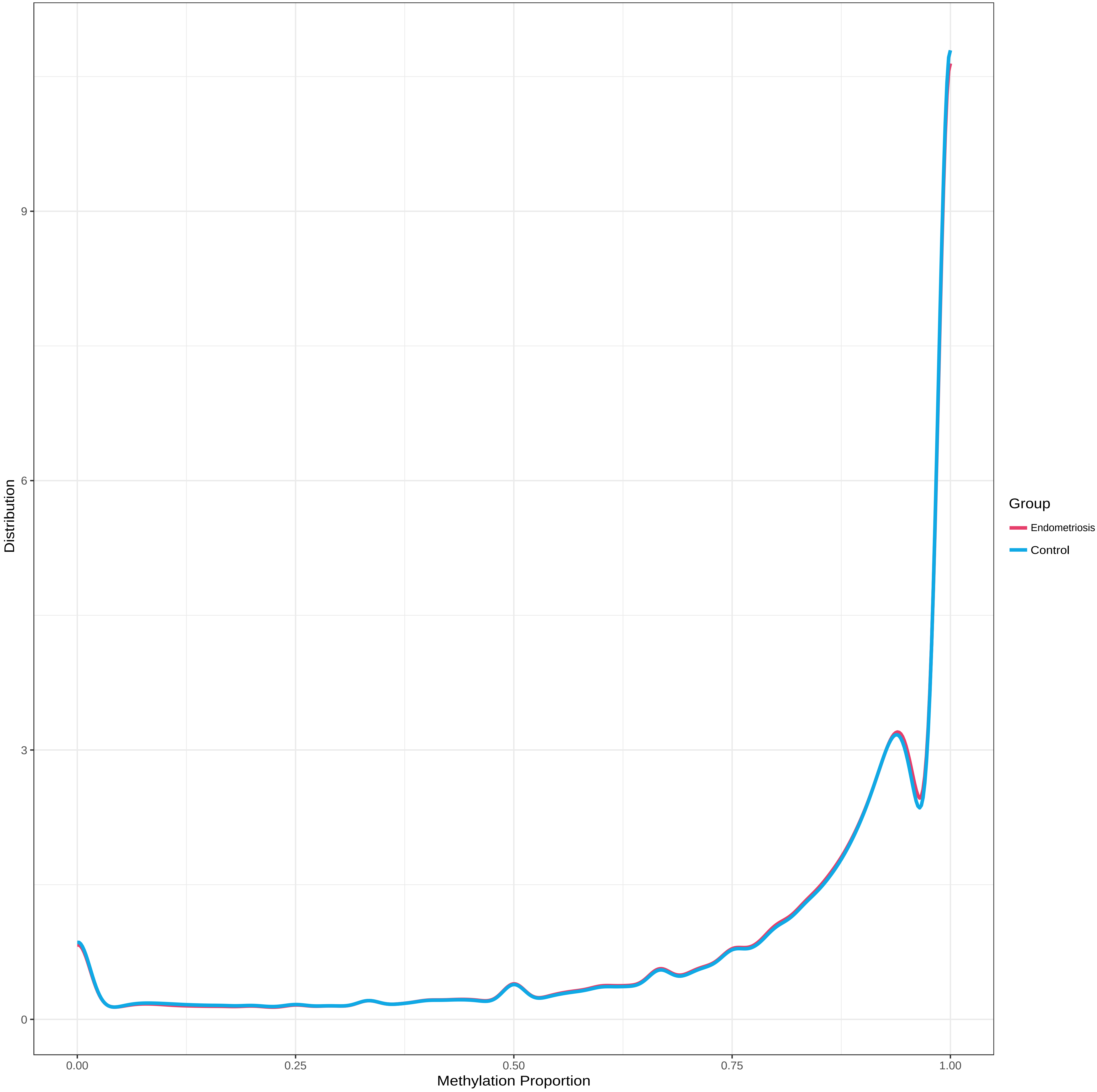


**Supplementary figure 1**: Methylation distribution across CpG sites and experimental groups.

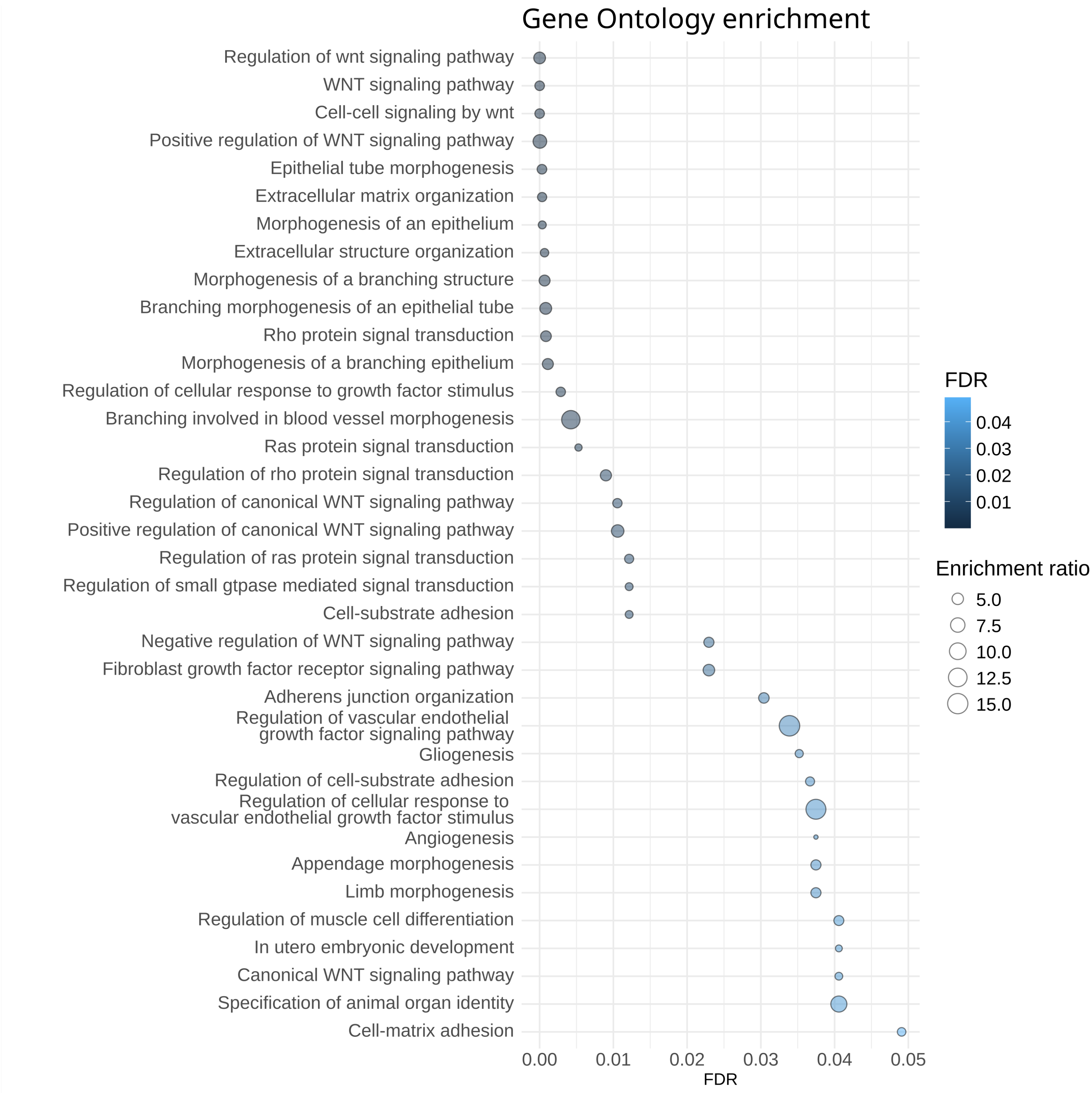


**Supplementary figure 2**: Dot plot summarizing the results of the ORA analysis for GO biological processes using genes associated with hypermethylated DMRs. Dot size represents the enrichment ratio, and colour intensity reflects the FDR values.

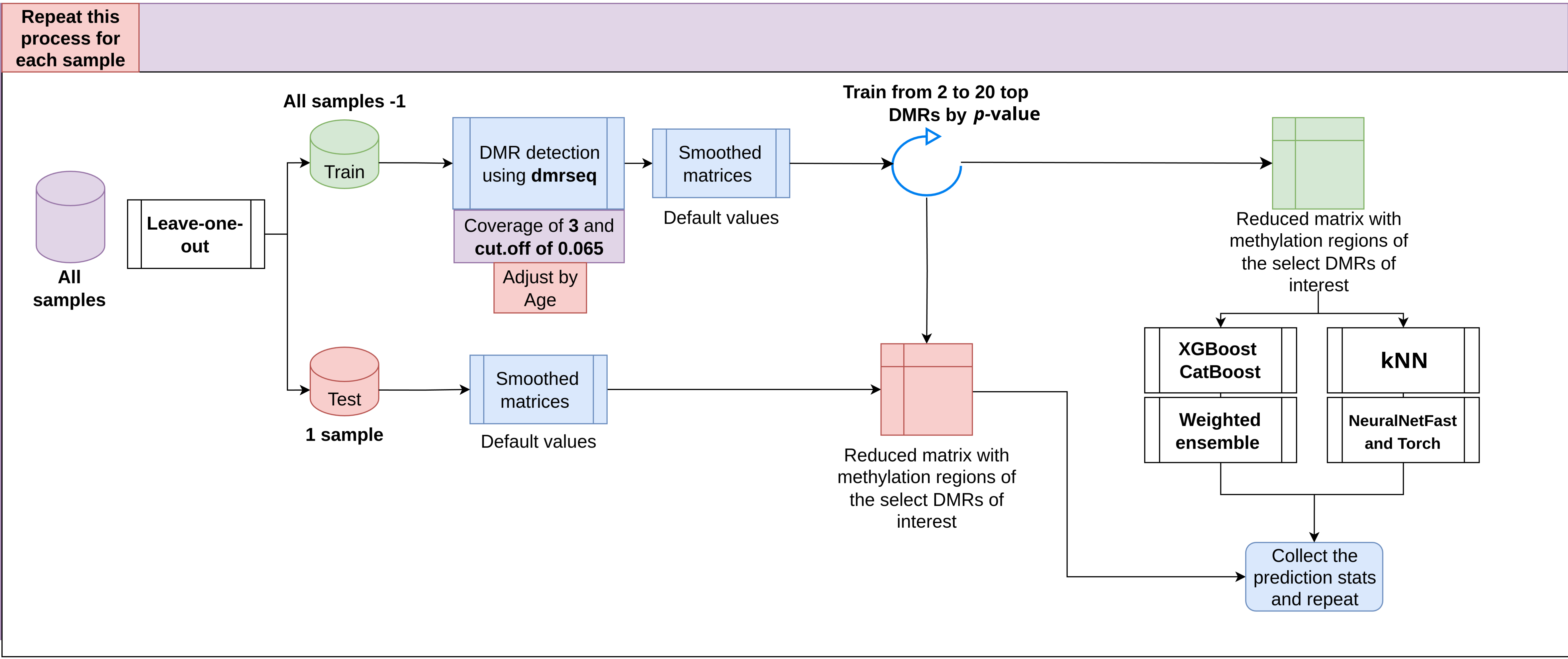


**Supplementary figure 3**: Diagram of the ML strategy and leave-one-out implementation.

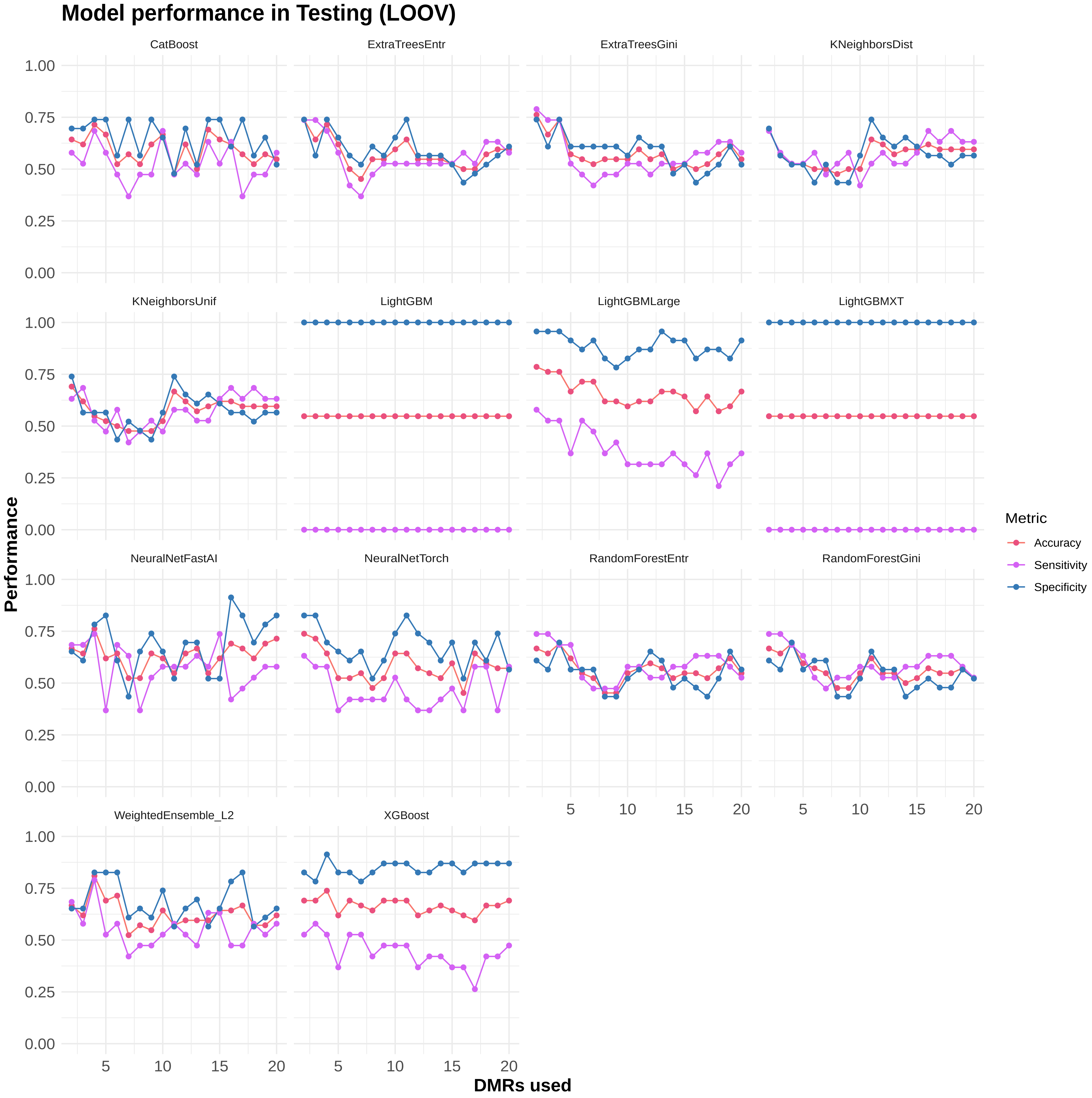


**Supplementary Figure 4**: Performance of the ML models in testing across different numbers of DMRs used.

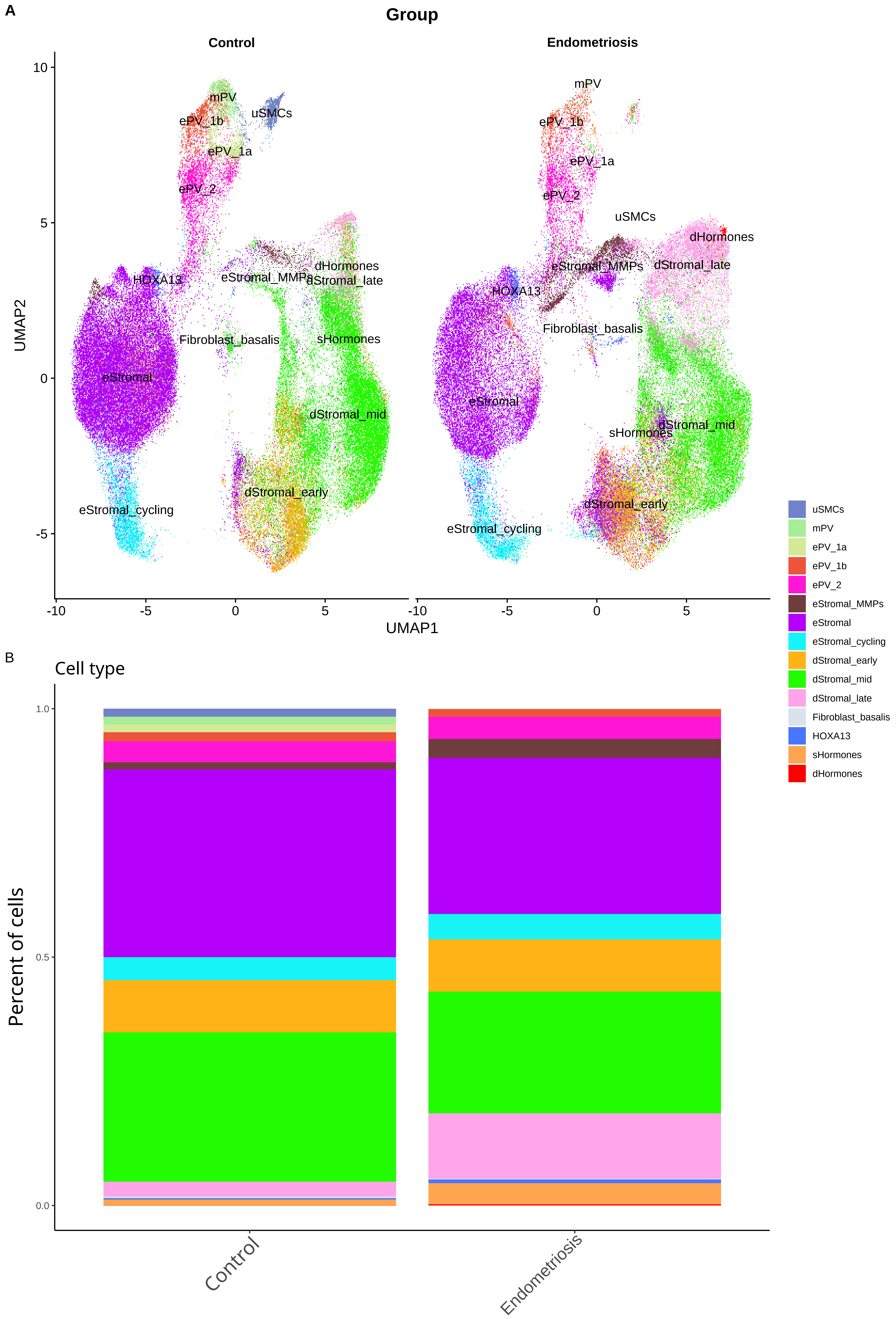


**Supplementary Figure 5**: (A) UMAP showing the different cell types within the mesenchymal lineage in control and endometriosis groups. (B) Bar plot illustrating the distribution of the detected cell types.
